# High-grade Ovarian Cancer Associated H/ACA snoRNAs Promote Cancer Cell Proliferation and Survival

**DOI:** 10.1101/2021.08.24.457387

**Authors:** Laurence Faucher-Giguère, Audrey Roy, Gabrielle Deschamps-Francoeur, Sonia Couture, Ryan M. Nottingham, Alan M. Lambowitz, Michelle S. Scott, Sherif Abou Elela

**Affiliations:** Département de microbiologie et d’infectiologie, Faculté de médecine et des sciences de la santé, Université de Sherbrooke, Sherbrooke, QC J1E 4K8, Canada; Département de biochimie et génomique fonctionnelle, Faculté de médecine et des sciences de la santé, Université de Sherbrooke, Sherbrooke, QC J1E 4K8, Canada; Institute for Cellular and Molecular Biology and Department of Molecular Biosciences and Oncology, University of Texas at Austin, Austin, Texas 78712, USA

## Abstract

Small nucleolar RNAs (snoRNAs) are an omnipresent class of non-coding RNAs involved in the modification and processing of ribosomal RNA (rRNA). As snoRNAs are required for ribosome production, the increase of which is a hallmark of cancer development, their expression would be expected to increase in proliferating cancer cells. However, the nature and extent of snoRNAs contribution to the biology of cancer cells remain largely unexplored. In this study, we examined the abundance patterns of snoRNA in high-grade serous ovarian carcinomas (HGSC) and serous borderline tumours (SBT) and identified a subset of snoRNA associated with increased invasiveness. This subgroup of snoRNA accurately discriminates between SBT and HGSC underlining their potential as biomarkers of tumour aggressiveness. Remarkably, knockdown of HGSC-associated H/ACA snoRNAs, but not their host genes, inhibits cell proliferation and induces apoptosis of model ovarian cancer cell lines. Wound healing and cell migration assays confirmed the requirement of these HGSC-associated snoRNA for cell invasion and increased tumour aggressiveness. Together our data indicate that H/ACA snoRNAs promote tumour aggressiveness through the induction of cell proliferation and resistance to apoptosis.

## Introduction

Ovarian cancer is the most lethal cancer of the female reproductive system (1). Most ovarian cancers are epithelial in origin, 70% being high grade serous carcinomas (HGSC) and 15% being serous borderline tumours (SBT) or tumours of low malignant potential (2,3). HGSC is aggressive and mostly diagnosed at a late stage, while SBT is normally much less aggressive and often diagnosed when still restricted to the ovaries. Currently, the diagnosis of SBT and its differentiation from HGSC relies on histological criteria including epithelial cell proliferation, stratified epithelium, microscopic papillary projections, cellular pleomorphism, nuclear atypia, mitotic activities and the absence of stromal invasion, which differentiates it from invasive carcinomas (3–6). Relying on histological criteria complicates early diagnosis and prevents definitive estimation of the invasive potential in the early stages of tumour development. In addition, microscopic analysis of tumours is not suitable for early detection and prediction of ovarian tumours.

Currently there are two tests being used for ovarian cancer screens, transvaginal ultrasound (TVUS) and measurement of blood levels of CA-125 protein (7–9). TVUS may only detect tumours of a certain mass and as such is not effective for very early stages of cancer development nor for differentiating between HGSC and SBT. On the other hand, CA-125 levels were found not to be useful for *de novo* screens since they are associated with a high rate of false positive results (10). Several, markers for ovarian cancer are being developed or considered but most of these markers are yet to be validated as accurate early predictors of tumour invasive potential. One common problem with these putative markers, which mostly takes the form of changes in the levels of protein coding mRNAs, is their reduced full-length extra-cellular stability preventing detection in patients’ fluids (1,11). Therefore, there is a need for identifying different types of markers that are more stable than mRNA and have the capability of predicting tumour potential.

Mid-size non-coding RNAs (mncRNAs) are the most abundant and stable class of non-ribosomal RNA in human cells, and are much easier to quantify using RT-qPCR than small non-coding RNAs like microRNAs (miRNAs) (12). mncRNA vary in size between 40-400 nucleotides. Furthermore, many of these mncRNAs are among the most stable RNAs in serum and patient’s fluids, like urine samples (13,14). This makes mncRNA a prime target for the identification of non-invasive biomarkers. Indeed, many types of cancer have been associated with small nucleolar RNAs (snoRNAs), which form the largest component of mncRNAs (12,15). snoRNAs are highly conserved and mostly function as a guide for the modification of ribosomal RNA (rRNA) and small nuclear RNA (snRNA). C/D box snoRNAs guide RNA 2’ O-ribose methylation, while H/ACA box snoRNAs guide pseudouridylation (16). These two classes of snoRNAs differ in their structure with the C/D box snoRNAs forming a loose stem loop, and the H/ACA box snoRNAs forming a tight, two-hairpin structure.

The highly structured nature of these two snoRNA classes increases their stability but at the same time makes their detection using standard sequencing methods difficult and unreliable (15,17). Recently, we have developed a mncRNA sensitive pipeline that accurately quantifies the different species of mncRNAs including snoRNAs (15). Thanks to the use of a highly processive thermostable bacterial reverse transcriptase (RT) and a quantification pipeline that recognizes multimapped and intron embedded sequencing reads, we are now able to directly quantify snoRNA abundance in cancer tissues using RNA-seq (15,18).

In this study, we have taken advantage of our newly developed mncRNA sensitive sequencing pipeline to identify stable RNA species that could differentiate between HGSC and SBT tissues and serve as potential markers for tumour aggressiveness. RNAs showing differential expression in SBT and HGSC were selected and validated using RT-qPCR and used to create a molecular HGSC signature. The relevance of HGSC-associated RNAs to the biology of ovarian cancer was examined by knocking them in using three different model ovarian cancer cell lines. Together our results identify three H/ACA snoRNAs as an effective HGSC signature and indicate that these H/ACA snoRNAs may promote tumour aggressiveness by preventing apoptosis and inducing cell proliferation and invasion potential.

## Materials and Methods

### Tissue sourcing RNA extraction, quality control and sequencing

HGSC and SBT tissues were obtained from the FRQS tissue bank (Université de Sherbrooke), the Ontario Tumour Bank (OTB) and the Alberta Research Tumor Bank (ARTB) (Table S1). Each 30 mg tissue sample was homogenized in 1 mL of TRIzol Reagent (Ambion) using a Polytron tissue homogenizer and the RNA extracted as previously described (19). RNA integrity was assessed using the 2100 Bioanalyzer (Agilent) and samples with RNA integrity index (RIN) > 4.8 were used for subsequent analyses. These values are available from - Table S1 for all samples. To further confirm the RNA integrity, we compared the expression levels of a panel of housekeeping genes (MRPL19, RPPH1, RNU6 and PUM1) to E. coli 23S rRNA spike-in (Roche, 10206938001) using RT-qPCR (20 picograms of E. coli ribosomal RNA was added to each sample during the RT preparation) and tissues exhibiting housekeeping levels within 2 standard deviations from the average relative expression were kept. The HGSC and SBT tissue identity was evaluated by comparing the expression level of VEGF-A and WFDC2 using RT-qPCR and HGSC tissues showing increased expression of VEGF-A or WFDC2 relative to SBT were used for sequencing or RT-qPCR. A cohort of randomly selected three HGSC and three SBT tissues were used for sequencing and the rest of tissues used for validation using RT-qPCR. TGIRT-seq libraries were constructed as previously described (15). ERCC spike-ins (20, 21) were added and used as control for detection uniformity.

### RNA sequence analysis

All datasets were analyzed using the same computational pipeline. Fastq files were checked for quality using FastQC v0.11.15 (20,21) and trimmed using Cutadapt v1.18 (22) and Trimmomatic v0.36 (23) (with TRAILING:30) to remove Illumina sequencing adapters and portions of reads of low quality, respectively. Reads were aligned using STAR v 2.6.1a (24) with the following options: --outFilterScoreMinOverLread 0.3, --outFilterMatchNminOverLread 0.3, --outFilterMultimapNmax 100, --winAnchorMultimapNmax 100, -- alignEndsProtrude 5 ConcordantPair. Only primary alignments were kept using samtools view -F 256 (v1.5) (25). The samples LGr RT157 and LGr RT35 librairies were split in multiple fastq files that were aligned separately then combined using samtools merge before quantification. The reference genome used was hg38 and the reference annotation to build the STAR alignment index was taken from Ensembl (v87) (26) supplemented with missing tRNAs and snoRNAs as described in (15). The gene quantification was done using CoCo correct_count module (v.0.2.0) (27) with the option -s 1 and -p. DESeq2 (28) was used for differential expression analysis using raw counts obtained by CoCo. As we performed non-fragmented sequencing, thus sequencing only small RNAs, equivalence between counts per million (CPM) and transcripts per million (TPM) can be assumed. Volcano plots were generated using R package Enhanced Volcano (EnhancedVolcano, RRID:SCR_018931, DOI:10.18129/B9.bioc.EnhancedVolcano). Heatmaps were generated using R package gplots. ROC curves were generated using the CombiROC online software (29). 3D structure of the ribosome with the different rRNA modifications was generated using the online database of the ribosomal modifications (3D Ribosomal Modification Maps Database, RRID:SCR_003097, (30).

### Cell culture and transfection

The ovarian adenocarcinoma SKOV3ip1 cell line (RRID:CVCL_0C84) was grown in DMEM/F12 (50/50) medium supplemented with 10% (v/v) fetal bovine serum (FBS) and 2 mM L-glutamine. TOV112D endometrioid carcinoma cells (ATCC Cat#CRL-11731, RRID:CVCL_3612) were grown in OSE medium supplemented with 10% (v/v) FBS and 2 mM L-glutamine. OVCAR-3 ovarian cancer cells (CLS Cat#300307/p690_NIH:Ovcar-3, RRID:CVCL_0465) were grown in RPMI 1640 supplemented with 20% FBS (v/v) FBS and 2 mM L-glutamine. Cell passaging was performed as recommended by the American Type Culture Collection (ATCC), no more than 20 passages were carried out for each cell lines. Each cell line was tested for mycoplasma contamination periodically (by qPCR method performed by the RNomics plateforme at the Université de Sherbrooke). Transfections were performed using Lipofectamine 2000 (Invitrogen) according to the manufacturer’s protocol with 15 nM of either siRNA or antisense oligonucleotide (ASO) (containing five 2’O methyl RNA bases, ten DNA bases with phosphorothioate bonds followed with five 2’O methyl RNA bases). The sequence of the different ASO and siRNA and equivalent scrambled controls are listed in Table S2. Cells were seeded at 5000 cells / well in 96-well plates and 300000 cells / well in 6-well plates. Plates were incubated for at least 48h at 37°C under 5% CO_2_.

### Cell line RNA extraction and quantification using RT-qPCR

RNA was extracted using the RNeasy mini kit (Qiagen), following the manufacturer’s guidelines with the on-column DNase digestion with the RNase-Free DNase set (Qiagen); except in this case, 1.5 volume of 100% ethanol was used to precipitate RNA in order to conserve small size RNAs. RNA quantification and integrity were assessed using the 2100 Bioanalyzer (Agilent). To carry out the reverse transcriptions, between 500ng and 1ug of RNA was used for cDNA synthesis using the MMULV-RT (Moloney Murine Leukemia Virus reverse transcriptase) (1 unit), RNaseOUT (20 units), dNTP (1mM) and random hexamers (3µM). The cDNA was diluted (3.33 ng / uL) and 10ng was used for the qPCR reactions. qPCR reactions were performed in 96-well plates using the Eppendorf Realplex Mastercycler ep gradient and in 384-well plates using the Bio-Rad CFX RealTime System. The reactions were performed in 10uL with 5uL Bio-Rad iTaq Universal SYBR Green Supermix, 3uL of diluted cDNA and specific primer pairs (200nM) (Table S3). No template and no RT reactions were used as negative controls. The normalized relative expression was calculated using the ΔΔCq where Cq is the quantification cycle. The statistical significance of the different conditions was calculated using a two-tailed Student’s t-test assuming unequal variance.

### Western blot

Cells were collected, pelleted and lysed in 50 µl lysis buffer (150 mM NaCl, 50 mM Tris-HCl (pH 8), 0.5 % NP40, 0.5 % Sodium Deoxycholate, 0.1 % Sodium Dodecyl Sulfate, 10 mM Sodium Pyrophosphate, and 50 mM EDTA) containing protease inhibitor cocktail (Complete, EDTA-free, Roche Diagnostics Indianapolis, IN) for 1 hour at 4°C in a tube rotator and centrifuged for 20 minutes. Protein concentration was determined using Bradford Protein Assay. Equivalent 10 µg protein samples were solubilized in Laemmli buffer, (62.5 mM Tris-HCl (pH 6.8), 10 % glycerol, 5 % β-mercaptoethanol and 2.3% SDS) denatured 3 minutes at 95°C and loaded onto 8% (EIF3A) and 12% (EIF4A2) gels. Electrotransfer to Hybond-ECL (GE Healthcare, Piscataway, NJ) was performed for 2 hours at 100 V. Membranes were blocked for 1 h at room temperature in TBS containing 0.1% Tween-20 and 5% skimmed milk. Membranes were hybridized overnight at 4°C in blocking solutions with agitation on a Nutator (Clay Adams Brand), with primary antibodies diluted at 1:1000 (Anti-EIF4A2: Abcam Cat#ab31218, RRID:AB_732123, and Anti-EIF3A: Abcam Cat#ab86146, RRID:AB_2096634) and 1:2000 (Anti-GAPDH: Novus Cat#NB300-221, RRID:AB_10077627). Membranes were washed in TBS containing 0.1% Tween-20. Secondary antibodies were diluted in blocking solution at 1:1000 (Anti-rabbit IgG: GE Healthcare Cat#NA934, RRID:AB_772206) and 1:2000 (Anti-mouse IgG: GE Healthcare Cat#NA931, RRID:AB_772210) and membranes were hybridized 90 minutes at room temperature with agitation on a Nutator (Clay Adams Brand). Membranes were washed and proteins were detected by chemiluminescence with Clarity Western ECL (Bio-Rad, Saint-Laurent, QC), using LAS 4000 (GE Healthcare, Mississauga, ON).

### Real time cell growth assay

XCELLigence (ACEA Biosciences Inc., San Diego, CA, USA) was used to analyze cell adherence in real time (31). E-plates were used to assess cell viability, while CIM plates were used to assess cell migration and invasion. For cell viability, 50μl of complete culture medium was added to each well of an E-plate. The plate was incubated for 30 minutes at 37°C with 5% CO_2_. Then, a concentration of 5000 cells per 50μl was added to each well. Measurements were taken every 10 minutes for three hours before the transfection. Following transfection, measurements were taken every 10 minutes up to at least 70 hours for a maximum of 90 hours. For cell migration assay, CIM plates were pre-assembled by adding 160uL of medium containing serum to each well of the bottom chamber of the CIM plate, the upper chamber was assembled on top and 50uL of serum free medium was added to each well. The plate was incubated for one-hour at 37°C with 5% CO_2_. The same procedure was used for cell invasion assay, except Matrigel gel (Corning, 356234) was added to coat the bottom of the upper chamber at a concentration of 1:40. Essentially as described before 50uL of Matrigel was diluted and added to the upper chamber and 30uL was removed to ensure proper coating of the chambers (31). The upper chamber was incubated for 5 h to ensure polymerization of the Matrigel. Bottom and upper chambers were assembled as mentioned previously. For both cell migration and invasion assay, cells were collected and 100ul were seeded at a concentration of 50000 cells/well (migration assay) or 25000 cells/well (invasion assay) 24 hours following transfection in 6-well plates. Measurements were taken every 10 minutes up to at least 70 hours for a maximum of 90 hours.

### Multiplex apoptosis and viability assay

The multiplex phenotypic assay was carried out as previously described (32). The medium was removed 48h after transfection in 96-well plates and 50µl coloration mix was added to each well. The coloration mix was composed of the four dyes at specific concentrations: Hoechst (Invitrogen), Calcein AM (with a 0.5 µM final concentration, Invitrogen), Alexa Fluor 647 (Invitrogen), Propidium Iodide (Invitrogen) in 4mL of Annexin V binding buffer (10mM HEPES, 140mM NaCl, 2mM CaCl2, pH 7.4). The plate was incubated for 30 minutes at 37 °C. The plate was then imaged using the Perkin Elmer Operetta. Nine pictures of different regions of each well were made and the images imported into the Columbus software (PerkinElmer, Waltham, MA, USA). This experiment was repeated four times to counteract any random biological variations.

### Cell proliferation assay

BrdU (5-Bromo-2-deoxyuridine) (Millipore, 203806) was used to assess cell proliferation and the assay was performed in 96-well plates, 48h after transfection. BrdU (0.75mM) was added to cells and incubated for 90 minutes at 37°C, followed by washing with PBS. To fix and permeabilize the cells, four different washing steps were conducted starting with paraformaldehyde (7.4%), then PBS, then Triton X-100 diluted in PBS (0.1%) and finishing with PBS. Bovine serum albumin 2% (BSA) in PBS was added followed by an incubation of 45 minutes at room temperature. DNase I (300μg/mL) was added, followed by an hour incubation at 37°C. The primary antibody, anti-BrdU (1:500) (Millipore Cat#MAB3222, RRID:AB_94758), was added and the plate incubated at room temperature for one hour. The cells were then washed twice with PBS, and the secondary antibody was added, goat anti-mouse IgG (H+L) Alexa fluor 488 (1:250 in 2% BSA) (Thermo Fisher Scientific Cat#A-11029, RRID:AB_2534088), as well as Hoechst stain (800ng/mL) (Thermo Fisher Scientific, Waltham, MA, USA). Cells were then incubated at room temperature in the dark for 45 minutes. A final wash of 100μL PBS was completed, and the plate was read with the Perkin Elmer Operetta. Nine pictures of different regions of each well were taken and the images imported into the Columbus software (PerkinElmer, Waltham, MA, USA).

### Flow cytometry assay

Flow cytometry assay was performed in 6-well plates. 48h after transfection, medium was removed and collected. Cells were PBS-washed, and trypsinized. Cells were collected and pelleted for 5 minutes at 100g, then suspended in ice-cold PBS. Cells were fixed by addition of cold ethanol (100% at -20°C), and then incubated at -20°C for a minimum of 30 minutes up to two weeks. Cells were again pelleted for 5 minutes at 400g, followed by 30 seconds at 16200g and washed in PBS. The double centrifugation and the PBS washing was repeated three times. RNase A (100μg/mL in PBS) was added, and the cells were incubated five minutes at room temperature. Propidium Iodide was then added, and the solution again incubated for 30 minutes at room temperature. Cells were then counted by FACS (Flow Cytometer BD Fortessa).

### Wound healing assay

Wound healing assays were performed 48h after transfection in 96-well plates. Culture medium was removed from the well and a scratch was performed in the middle of the well with a 200uL tip. Medium was added containing hydroxyurea (60mM) to inhibit cell proliferation. Using the Cloneselect apparatus (Molecular Devices) pictures of each well were taken 24h, 48h and 72h after the scratch.

## Results

### Identification of high-grade ovarian cancer associated mid-size non-coding RNA

To identify potential biomarkers of tumour aggressiveness, we compared the expression pattern of mid-size non-coding RNAs in HGSC and SBT, which are similarly composed of mostly epithelial cells, thus minimize the risk of cell type bias (33). The screening process was performed in two stages: first, to discover HGSC-associated mncRNAs and second, to validate the association (Figure 1A). In the discovery phase we used our recently developed mid-size non-coding RNA sensitive sequencing method (non fragmented TGIRT-seq) to identify RNAs that are differentially expressed in HGSC. Subsequently, the association of the RNA expression with HGSC was confirmed using RT-qPCR in the validation phase. The discovery screen was performed using 3 SBT and 3 HGSC tissues, while the validation screen was performed using an independent set of 18 SBT and 19 HGSC samples. All extracted RNA had an RNA integrity number (RIN) > 4.8 and resulted in a stable and consistent RT-qPCR amplification of four different housekeeping genes (MRPL19, RPPH1, RNU6 and PUM1) confirming the quality and integrity of the RNA. The identity of the selected tissues was confirmed both by the pathology report and the expression levels of VEGF-A (34–36) and WFDC2 (37,38), which were shown to be upregulated in HGSC.

**Figure 1.**
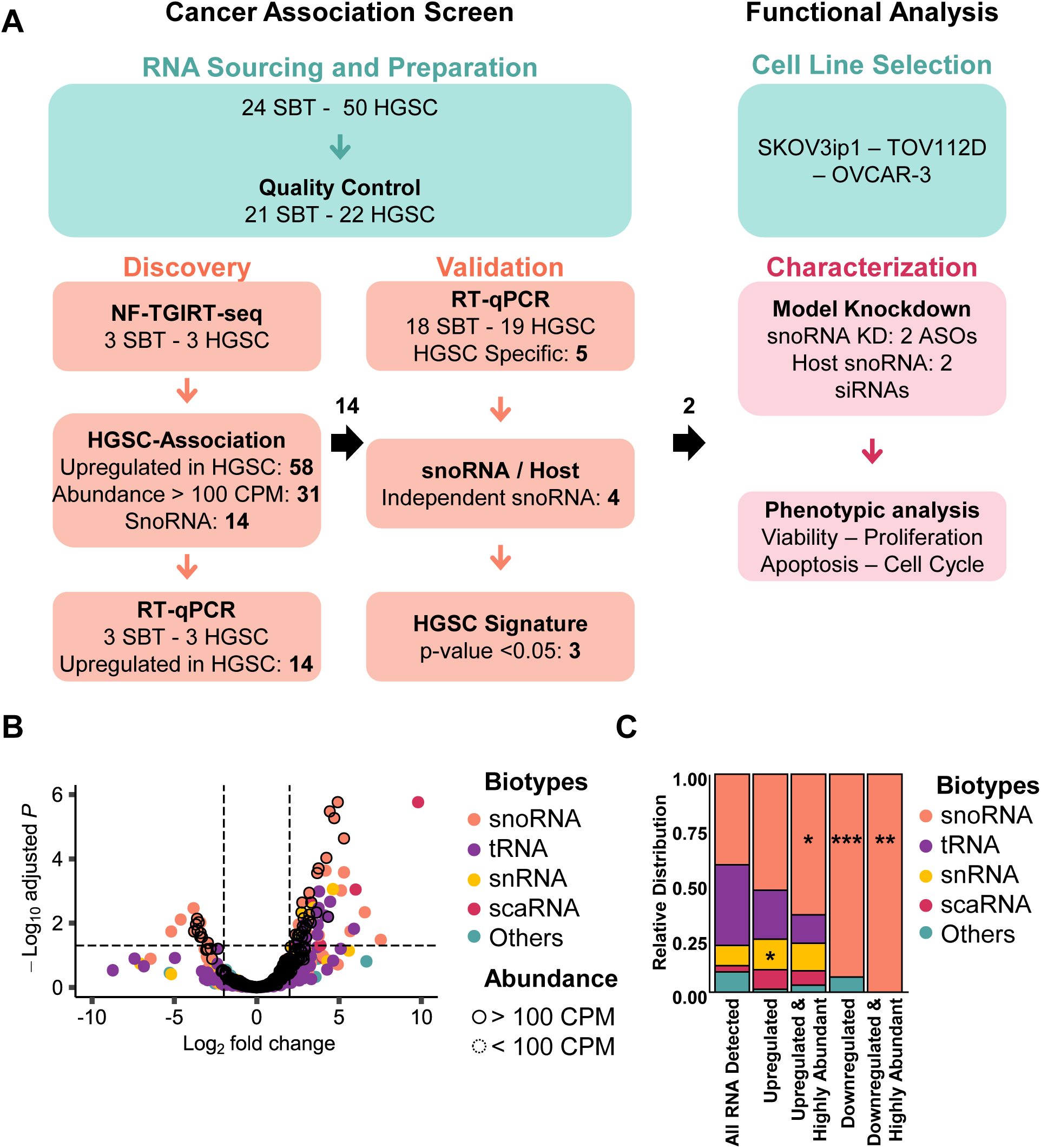
snoRNA expression is specifically modified in high-grade ovarian cancer. **(A)** Strategy for the identification of high-grade ovarian cancer associated mid-size non-coding RNA. RNA was extracted from serous borderline tumours (SBT) or high-grade serous ovarian cancers (HGSC) and the RNA and tissue quality (QC) were assessed using RT-qPCR. Total non-fragmented RNA was sequenced using TGIRT-seq (NF-TGIRT-seq) and mid-size non-coding RNA (mncRNAs) ranging in size between 40-400 nucleotides and generating more than 100 counts per million (CPM) were considered for association with HGSC. RNAs that are upregulated by more than 2-fold in HGSC and with a Benjamini-Hochberg adjusted P-value ≤ 0.05 were verified using RT-qPCR in the same sequenced tissues. snoRNAs that were upregulated in HGSC independent of their host genes were knocked down using antisense oligonucleotides (ASO) and evaluated for function using three model ovarian cancer cell lines. **(B)** snoRNAs are the most abundant class of HGSC-associated mncRNAs. The log2 fold change in abundance of HGSC RNA relative to SBT RNA as measured by NF-TGIRT-seq was plotted against the –log10 adjusted p-value. The most significantly down- and up-regulated mncRNAs in HGSC are in the upper most left and right quadrant of the volcano plot, respectively. Dots highlighted with a dark contour indicate RNA expressed more than 100 CPM. The name of the different RNA classes is indicated on the right. **(C)** snoRNAs are specifically dysregulated in HGSC. The relative distribution of RNAs that are detected (>1 CPM), differentially expressed (- or + log2 fold change and adjusted p-value of at most 0.05), and highly abundant (>100 CPM) in HGSC. The name of the different RNA classes is indicated on the right. The stars (*, ** and ***) indicate the significance as measured by Chi-square tests with p-values of 0.03, 0.02, 0.001 and 0.0003 respectively when comparing the relative enriched proportion of each mncRNAs between the full dataset (All RNA Detected) and the subsets of interest.

As indicated in Figure 1B, all four main biotypes of mncRNA (snoRNA, tRNA, snRNA and scaRNA) were detected by sequencing and the majority were similarly expressed in both SBT and HGSC tissues. This suggests that the overall expression program of mncRNA is not modified in HGSC when compared to SBT. Instead, a small subset of 58 mncRNAs present at >1 CPM were upregulated in HGSC with a log2 fold change of >2 and a Benjamini-Hochberg adjusted p-value of at most 0.05 when compared to the RNA extracted from SBT (Figure 1B). The majority of the RNAs that were differently expressed in HGSC were tRNAs or snoRNAs. From these, most of the differentially expressed RNAs present at >100 CPM were snoRNAs (Figure 1C). Indeed, the results indicated that in HGSC samples, 65% of all highly expressed mncRNAs that were upregulated were snoRNAs as well as 100% of all highly expressed mncRNA genes that were downregulated. Strikingly, snoRNAs were the only biotype showing statistically significant enrichment in abundance proportion between all RNA detected and the dysregulated (either up- or downregulated) and highly abundant categories between HGSC and SBT (Figure 1C). These data indicate that the expression of snoRNAs is specifically modulated in HGSC.

### H/ACA and C/D box snoRNAs are differentially regulated in high-grade ovarian cancer

The observed changes in the abundance of snoRNAs between the two tissue types may stem from a general dysregulation of the ribosome biogenesis machinery or gene-specific modulation of gene expression. To differentiate between these two possibilities, we compared the abundance of H/ACA and C/D snoRNAs in HGSC and SBT tissues. Surprisingly, the results indicated that H/ACA and C/D snoRNAs are inversely regulated in HGSC tissues (Figure 2A and B). The overall expression of H/ACA snoRNAs tended to increase in HGSC, while the expression of C/D snoRNAs tended to decrease (Figure 2A). Analysis of the amplitude and statistical significance of the change in the abundance of snoRNAs indicated that, while a small subset of C/D snoRNAs were significantly downregulated most of the H/ACA snoRNAs were upregulated in HGSC (Figure 2B). Indeed, no H/ACA snoRNA was downregulated in HGSC and only two C/D snoRNAs were upregulated and highly expressed in HGSC (Figure 2C). The 9 downregulated C/D snoRNAs represented 3.6% of the expressed C/D snoRNAs and were all part of the highly duplicated SNORD113-SNORD114 family embedded within the intron of lncRNA MEG8, which was previously linked to pancreatic and lung cancer epithelial to mesenchymal transition (EMT) (39,40). The two C/D snoRNAs that were highly expressed and upregulated in HGSC (SNORD72 and SNORD101) represented only 0.8% of all expressed C/D snoRNAs and were embedded in two different ribosomal protein host genes (RPL37 and RPS12). This indicates that high-grade ovarian cancer associates with changes in the expression of a small subset of C/D snoRNAs that are mostly more highly expressed in borderline tumours.

**Figure 2.**
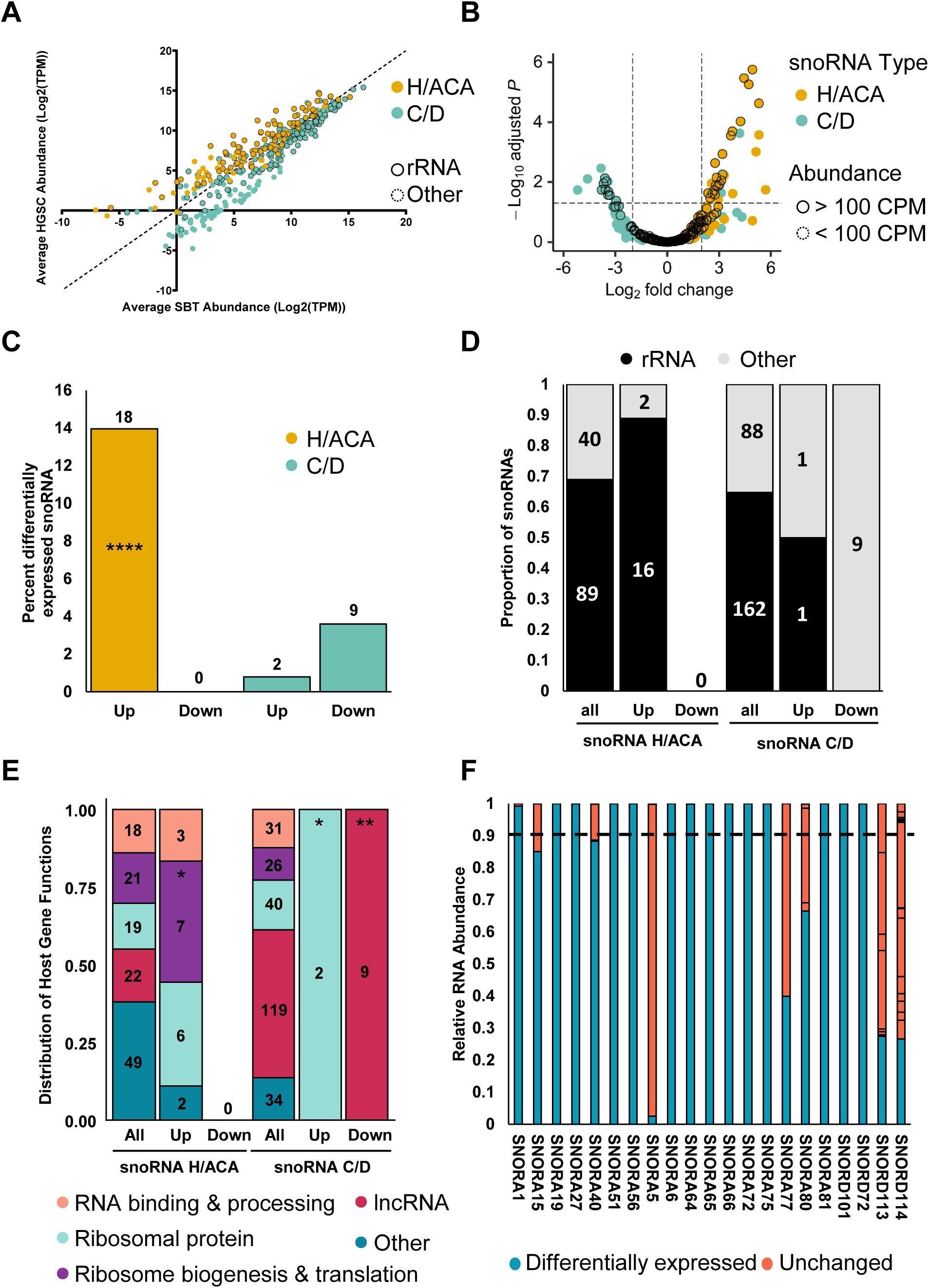
H/ACA and C/D box snoRNAs are differentially expressed in HGSC. **(A)** H/ACA and C/D box snoRNAs display distinct abundance patterns in HGSC and SBT. The abundance of H/ACA and C/D snoRNAs as determined by TGIRT-seq was compared using a scatter plot. C/D snoRNAs are shown in cyan and H/ACA snoRNAs are shown in yellow for panels A-C. Dots highlighted with a dark contour indicate snoRNAs with predicted rRNA modifications targets. **(B)** H/ACA snoRNAs are the most upregulated class of snoRNA in HGSC. The log2 fold change of snoRNA abundance in HGSC relative to that of SBT was plotted against the –log10 adjusted p-value. The most significantly down- and up-regulated mncRNAs in HGSC are in the upper most left and right quadrant of the volcano plot, respectively. Highlighted dots indicate RNAs expressed above 100 CPM. **(C)** C/D snoRNAs are mostly down regulated in HGSC. Bar graph showing the percent of snoRNAs that are highly expressed (expressed above 100 CPM) and either up or down regulated in HGSC. The number above the bars indicate the number of snoRNAs that fall within that category. The proportion of upregulated H/ACA snoRNAs is significantly higher than the proportion of C/D snoRNAs in HGSC (Fisher’s exact test ****p < 0.0001). **(D)** Most HGSC-associated snoRNAs target ribosomal RNA modifications. The proportion of snoRNAs that target rRNA modifications is indicated for both up and down regulated snoRNAs in the form of stacked bar graph. The number inside the bars indicate the number of snoRNAs that fall within that category. All, up and down indicate respectively all detected snoRNAs (at least 1 CPM in one or more tissue), upregulated or downregulated in HGSC. **(E)** Most upregulated snoRNAs are expressed from host genes coding for proteins associated with ribosome biogenesis and translation. The function of the host genes harboring all detected, upregulated, and downregulated snoRNAs is indicated in the form of a stacked bar graph. The categories of the host gene functions are indicated on the bottom. The proportion of snoRNAs encoded in host genes with specific functions were compared between all detected snoRNAs and those in subgroups of interest, per snoRNA class. The star (*) indicates p-values ranging from 0.05 to 0.03 while ** indicate a p-value of 0.006. **(F)** Most HGSC-associated snoRNAs are expressed from a single expressed gene copy. The relative abundance of the RNA generated from the different gene copies coding for each of HGSC-associated snoRNA is indicated in the form of a bar graph. The differentially expressed snoRNA copies are indicated in blue and those produced from other copies indicated in pink. The dashed line is the cut-off at 90%.

Unlike C/D snoRNAs, none of the H/ACA snoRNAs were downregulated in HGSC and 14% of all expressed H/ACA snoRNAs (18 snoRNAs) were upregulated in HGSC (Figure 2C). The higher proportion of upregulated H/ACA in HGSC compared to C/D was found to be statistically significant (Fisher’s exact test, p < 0.001). Interestingly, the majority (16/18) of upregulated H/ACA snoRNAs are predicted to guide rRNA modifications, indicating that the upregulation of these snoRNAs may signify changes in ribosome biogenesis and/or function (Figure 2D). Strikingly, the opposite trend was found for the C/D snoRNAs with dysregulated expression in HGSC (either up or down regulated), as the majority (9/10) were not predicted to guide rRNA modifications, suggesting different roles for the two snoRNA classes in ovarian cancer tumorigenesis. Consistent with the findings for HGSC upregulated H/ACA snoRNAs, a significant proportion (13/18) are embedded within ribosome related genes, most specifically genes which encode ribosomal proteins, or proteins involved in ribosome biogenesis and translation (Figure 2E). Many snoRNAs are known to be produced from multicopied genes (41) and thus more than one gene can generate the exact same snoRNA or a snoRNA with very close sequence. Such snoRNAs with very close sequence are referred to as snoRNAs of the same family. Care must thus be taken when quantifying same-family snoRNAs using RNA-seq and qPCR. Interestingly, most snoRNAs showing modified expression in HGSC were primarily generated from a single expressed gene, except for the highly duplicated SNORA5, SNORD113 and SNORD114 (Figure 2F). Of the 20 upregulated snoRNAs (2 C/D and 18 H/ACA snoRNAs), 14 snoRNAs represented at least 90% of the sum of the abundance of all copies of their family (2 C/D and 12 H/ACA snoRNAs). These 14 mostly monogenic snoRNAs are more likely to be homogeneously regulated in HGSC and thus were selected for further study.

### Identification of HGSC snoRNA signatures

To validate the predicted upregulation of the 14 abundant snoRNAs in HGSC, we monitored their expression using RT-qPCR in the same tissues used for sequencing. As indicated in supplementary Figure 1, the sequencing and RT-qPCR expression values are well correlated supporting the conclusion of the RNA-seq. To confirm the association of the snoRNAs with HGSC and evaluate their potential as possible biomarkers, we monitored their expression using RT-qPCR in an independent cohort of 19 HGSC and 18 SBT tissues. As indicated, in Figure 3A, five snoRNAs (SNORA81, SNORA56, SNORD72, SNORA6 and SNORA19) were mostly uniformly overexpressed in HGSC when compared with SBT and displayed similar or better association with high-grade ovarian cancer than the known mRNA markers VEGF-A and WFDC2. The expression of these five snoRNAs was significantly upregulated in all HGSC tissues tested as compared to SBT tissues with Benjamini-Hochberg adjusted p-values between 0.002 and 0.04 (Figure 3B).

**Figure 3.**
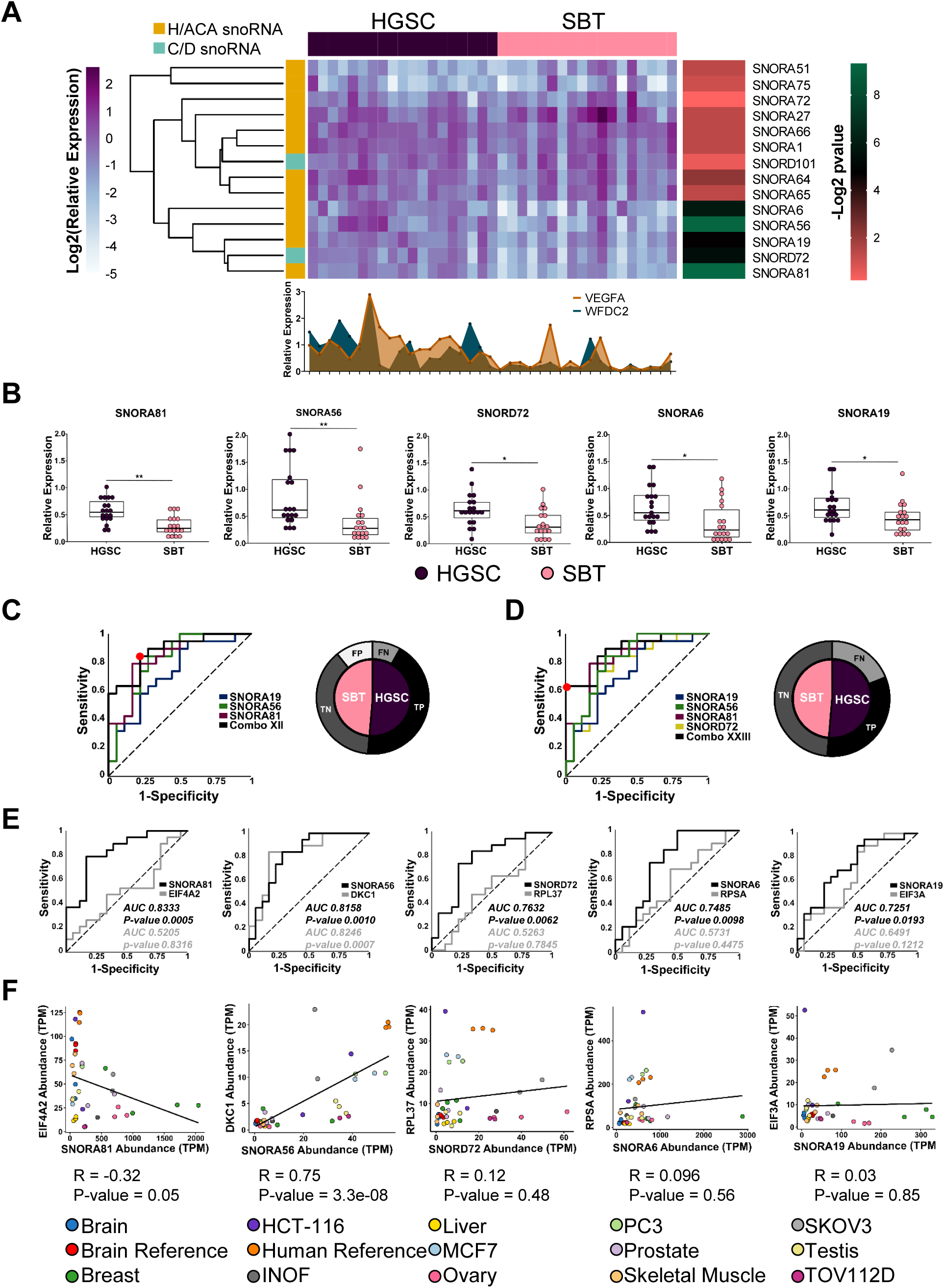
Identification of a HGSC snoRNA signature. **(A)** Five snoRNAs accurately discriminate between HGSC and SBT derived from independent patients’ cohorts. The relative abundances of the 14 HGSC-associated snoRNAs identified by sequencing were examined in a blind mix of 37 tissues using RT-qPCR. Data were normalized relative to the mean amount of MRPL19 mRNA and a spike-in of *E. coli* 23S rRNA, and the log2 of the relative expression is presented in the form of a horizontally clustered heatmap. The snoRNA class is indicated on the left, the identity of the tissue of origin shown on the top and the significance (-log2 of Benjamini-Hochberg adjusted p-value) of differential expression between HGSC and SBT is shown on the right. For comparison, the histogram shown at the bottom indicates the relative expression of two known HGSC molecular markers (VEGF-A and WFDC2). **(B)** Comparison between the distributions of the abundance of the top five HGSC-associated snoRNAs in 18 SBT and 19 HGSC tissues. The abundance of the snoRNAs was determined as described in A and the data shown as a box plot. Each point in the box plot represents one tissue and the data obtained from HGSC and SBT are shown in purple and pink, respectively. The stars indicate Benjamini-Hochberg adjusted p-values from left to right of 0.002 (**), 0.002 (**), 0.02 (*), 0.02 (*) and 0.04 (*). **(C & D)** Identification of optimal snoRNA signature of HGSC. Thirty-one different combinations of the five HGSC-associated snoRNAs were evaluated and the receiver operating characteristic (ROC) curve of the most sensitive (C) and specific (D) combinations are shown. The curves show the snoRNA combinations in black (Combo XII and Combo XXIII). The optimal discrimination cut-off is indicated by the red dot. The pie charts shown on the right indicate the signature dependent distribution of the 37 tissues examined. SBT and HGSC are indicated in pink and purple, respectively. Tissues that are correctly identified are indicated in black and dark grey, while tissues that are misidentified are indicated in light grey and white, respectively. **(E)** snoRNAs may discriminate between HGSC and SBT in a host gene independent manner. The capacity of each of the top five HGSC-associated snoRNAs to discriminate between HGSC and SBT tissues was compared to that of their host genes using ROC curves. Confidence intervals were calculated using the Wilson/Brown method (57). Black and grey lines indicate the ROC curve of the snoRNAs and host genes, respectively, while the dotted line indicates a random distribution. The area under the curve is indicated for each snoRNA (black) and each host gene (grey) with the respective p-value associated with the confidence intervals, calculated using the Wilson/Brown method. **(F)** The expression of all but one HGSC-associated snoRNAs and their host genes is independently regulated. The abundance of the top 5 HGSC-associated snoRNAs and their host mRNAs in 7 healthy tissues and six cell lines was retrieved from published TGIRT-seq datasets (39,42) and presented as a scatter plot. The Pearson correlation for the different gene pairs is indicated below each graph.

To evaluate the potential role as biomarkers for these snoRNAs, we evaluated whether the combination of multiple snoRNAs abundance would increase the sensitivity to discriminate between HSGC and SBT. To identify the optimal HGSC signature that best discriminates between HGSC and SBT with the highest sensitivity and accuracy, we evaluated the discrimination value of different snoRNA combinations. We examined 31 different combinations of snoRNAs (all combinations of 1, 2, 3, 4 and 5 snoRNAs amongst this group) resulting in the identification of SNORA56, SNORA19 and SNORA81 as the best HGSC signature with a maximum sensitivity value of 0.8 and specificity of value of 0.25 (Figure 3C). The second-best combination included four snoRNAs (SNORA81, SNORD72, SNORA56 and SNORA19) with sensitivity and specificity values of 0.6 and 0.0 respectively. At the optimal cut-off, the three snoRNAs signature accurately identified most HGSC tissues with four false positives and three false negatives. In contrast, the four snoRNA signature resulted in zero false positives and seven false negatives at the optimal cut-off. Furthermore, the three snoRNAs signature could significantly discriminate (Fisher’s exact test *** p-value 0.0002), through a clustered heatmap, between the blind mix of 37 HGSC and SBT tissues (supplementary Figure 2). We conclude that SNORA56, SNORA19 and SNORA81 form the optimal H/ACA snoRNAs signature for the identification of HGSC tissues.

### Induction of snoRNA expression in HGSC is not linked to host gene expression

The newly identified HGSC-associated snoRNAs are all expressed as part of introns of protein coding genes. This raises the question of whether the association of these snoRNAs with HGSC signifies a direct link between snoRNAs and the tissue type or an indirect effect of HGSC dependent induction of their host gene. To discriminate between these two possibilities, we compared the expression of the five HGSC-associated snoRNAs with the expression of their host genes in HGSC and SBT tissues. As indicated in Figure 3E, in all cases except that of SNORA56, the snoRNAs discriminated between HGSC and SBT independently of their host gene expression. In the case of SNORA56, both snoRNA and host gene *DKC1*, which encodes the H/ACA snoRNA binding protein and pseudouridylase dyskerin, were able to discriminate between SBT and HGSC tissues. This is likely due to the shared function of H/ACA snoRNAs and this interacting core protein. This indicates that in most cases the abundance of snoRNAs in cells is not linked to the amount of the host gene mRNA. To evaluate the validity of this assumption we compared the expression levels of the HGSC-associated snoRNAs and their host genes in different unrelated tissues and cell lines using previously published TGIRT-seq datasets (39,42). As indicated in Figure 3F, we found no correlation between the snoRNA and host gene mRNA abundance except in the case of SNORA56 / DKC1 gene pair. Once again, these data indicate that the expression of snoRNA and host gene is not strictly linked but depends mostly on the function of the host gene. We conclude that factors independent of host gene expression contribute to the increased abundance of a subset of H/ACA snoRNAs in HGSC.

### Knockdowns of high-grade ovarian cancer associated snoRNAs inhibit cell proliferation

To evaluate the biological significance of the HGSC-associated snoRNAs, we knocked down SNORA19 and SNORA81 in the model ovarian cancer cell lines SKOV3ip1, TOV112D and OVCAR-3. Among those, we chose SNORA19 and SNORA81 as the models that represent snoRNAs with the lowest and highest degrees of association with HGSC (Figure 3E). These two snoRNAs are embedded in the introns of the genes encoding eukaryote translation initiation factors EIF3A and EIF4A2 (Figure 4A). EIF3A introns harbour a single copy of SNORA19, while EIF4A2 introns contain SNORA81 along with four other non-HGSC-associated snoRNAs (SNORA4, SNORA63, SNORA63B and SNORD2). To differentiate between snoRNA and host gene effects, we knocked down the snoRNAs using two independent antisense oligonucleotides (ASOs) which target the RNA for degradation using RNase H and compared the results to the knockdown of host genes using two independent siRNAs targeting a mature RNA splice junction (Figures 4A and Supplementary Figure 3A). The ASOs target both mature snoRNA and the unspliced pre-mRNA and as such are expected to reduce the amount of both host mRNA and snoRNA levels. In contrast, the siRNAs specifically target the spliced sequence and as such should not affect the accumulation of snoRNA. As predicted, the ASOs targeting SNORA19 and SNORA81 reduced the abundance of both snoRNA and host mRNA, while the siRNAs targeting EIF3A and EIF4A2 decreased the abundance of the host mRNA without affecting the embedded snoRNAs (Supplementary Figure 3B and C). Examination of the effect of the knockdown on the level of EIF3A and EIF4A2 proteins indicated that as expected the siRNAs against the mature RNA reduce the level of the protein in the cell. Only the ASOs against SNORA19 and not those against SNORA81 reduced the level of the protein produced by the host gene (Supplementary Figure 3D and E). Similar knockdown effects were observed in all three ovarian cancer cell lines tested (Supplementary Figure 4). This indicates that SNORA81 could be knocked down with little or no effect on the host gene proteins, while knockdown of SNORA19 inhibits both snoRNA and host RNA and protein expression.

**Figure 4.**
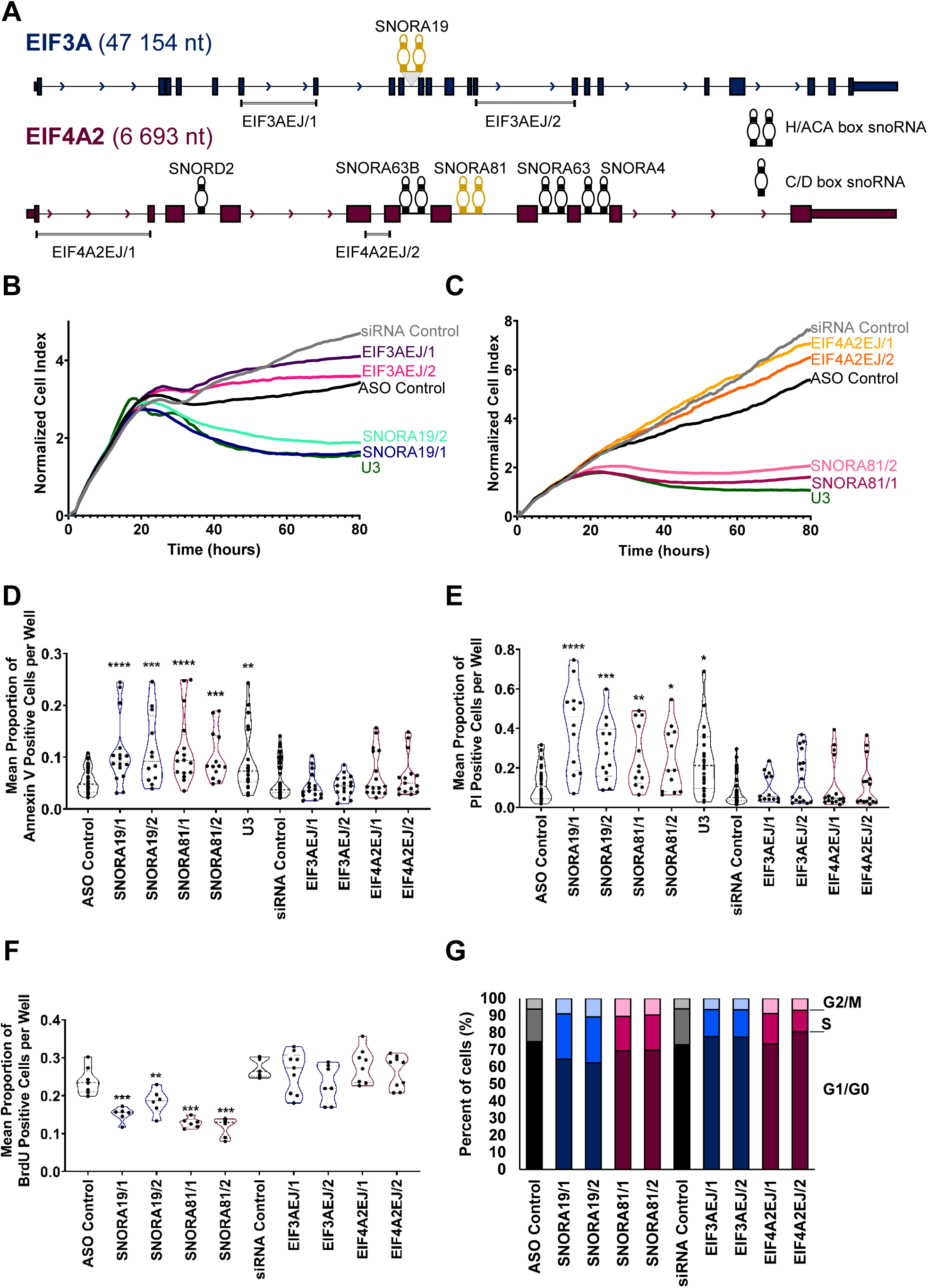
HGSC-associated snoRNAs induce cell survival and inhibit apoptosis. **(A)** SNORA19 and SNORA81 are expressed from host genes encoding the translation initiation factors EIF3A and EIF4A2 respectively. The structure of the two snoRNA encoding genes is depicted in scale. Exons, introns, and snoRNAs are indicated in black boxes, lines and stem-loops, respectively. Both SNORA19 and SNORA81 are indicated in yellow. The names of the snoRNAs are indicated above them and the position of the exon-junction siRNAs used to inhibit the expression of the mature mRNA are shown at bottom. Arrows indicate the direction of transcription and gene sizes are shown on the left. **(B)** Knockdown of SNORA19 represses cell growth. The ovarian cancer model cell line SKOV3ip1 was transfected with two ASOs against SNORA19 (SNORA19/1 and SNORA19/2) or two siRNAs targeting the exon junction of EIF3A mRNA (EIF3AEJ/1 and EIF3AEJ/2) and the effect on growth was followed in real-time through changes in cell impedance. Unrelated siRNAs and ASOs were used as negative controls and an ASO against U3 snoRNA was used as a positive control. The y-axis indicate the normalized cell index, which is the average of the three technical replicates. The experiment was carried out in three biological replicates. **(C)** Knockdown of SNORA81 represses cell growth. Depletion of SNORA81 (SNORA81/1 and SNORA81/2) and its host gene (EIF4A2EJ/1 and EIF4A2EJ/2) was carried out as described in B. The y-axis indicate the normalized cell index, which is the average of the three technical replicates. The experiment was carried out in three biological replicates. **(D)** Knockdowns of SNORA19 and SNORA81 induce apoptosis. The effect of the different knockdowns described in B and C on apoptosis was evaluated 48 hours after transfection using Annexin V assays. The results, shown as violin plots, represent the mean proportion of annexin V positive cells per well. Mann-Whitney tests were performed and are indicated by asterisks. The stars indicate p-values from left to right for SNORA19/1 (p-value <0.0001, ****), SNORA19/2 (p-value 0.0009, ***), SNORA81/1 (p-value <0.0001, ****), SNORA81/2 (p-value 0.0002, ***) and U3 (p-value 0.008, **), as compared to the control ASO. The experiment was carried out in three biological replicates with at least four technical replicates. **(E)** Knockdown of SNORA19 and SNORA81 induce necrosis. The experiment was carried out as in D, but necrosis caused by contact of Propidium iodide and nucleic acids in cells with compromised membranes was assessed. The stars indicate p-values from left to right for SNORA19/1 (p-value <0.0001, ****), SNORA19/2 (p-value 0.0004, ***), SNORA81/1 (p-value 0.01, **), SNORA81/2 (p-value 0.04, *) and U3 (p-value 0.01, *) as compared to the control ASO. The experiment was carried out in three biological replicates with at least four technical replicates. **(F)** Knockdowns of both SNORA19 and SNORA81 inhibit cell proliferation. SKOV3ip1 were transfected as described in B and C and labelled with BrdU 48 hours after transfection. The violin plots show the mean proportion of BrdU positive cells per well. Mann-Whitney tests were performed, and significance is indicated by asterisks. The stars indicate p-values from left to right for SNORA19/1 (p-value 0.0007, ****), SNORA19/2 (p-value 0.008, **), SNORA81/1 (p-value 0.0007, ***), SNORA81/2 (p-value 0.0007, ***) as compared to the control ASO. The experiment was carried out in three biological replicates. The experiment was carried out in three biological replicates with at least four technical replicates. **(G)** Knockdown of SNORA19 and SNORA81 causes arrest in the S phase of the cell cycle. SKOV3ip1 cells transfected and propidium iodide-stained were analyzed by FACS to determine their cell cycle phase after 48h and these proportions are displayed as stacked bar graphs. The cell cycle phases detected after the knockdown of SNORA19 and SNORA81 are shown in shades of blue and purple, respectively. The experiment was carried out in three biological replicates.

To assess the contribution of HGSC-associated snoRNAs to cell growth, we compared the cell growth pattern of SKOV3ip1 cell line in real time before and after knockdown of the snoRNAs and the host genes. As indicated in Figure 4B and C, the knockdown of SNORA19 and SNORA81, and not the of the host genes, strongly inhibited cell growth. Strikingly, the inhibition of SNORA19 and SNORA81 inhibited cell growth to the same levels observed after the knockdown of the highly conserved U3 snoRNA essential for rRNA processing and ribosome production (43). This clearly indicates that SNORA19 and SNORA81 and not their host genes are required for the growth of cancer cells. The snoRNA dependent growth is likely due to increased cell survival and increased proliferation rate. As indicated in Figure 4D and E, the knockdown of the snoRNAs and not of their host genes increased apoptosis and necrosis. In addition, the knockdown of the snoRNAs decreased cell proliferation and delayed the S phase of the cell cycle (Figure 4F and G). The effects of SNORA81 and SNORA19 on apoptosis, necrosis, proliferation, and cell cycle were observed in all three different cell lines tested (Supplementary Figure 5). The results were similar to the effect observed with U3 knockdown, indicating a major effect on ribosome biogenesis and/or function. We conclude that these HGSC-associated snoRNAs may increase tumour aggressiveness by promoting cell proliferation and resistance to apoptosis.

To better assess the contribution of HGSC-associated snoRNAs to cancer cell aggressiveness, we examined the impact of SNORA19 and SNORA81 on migration, wound healing and invasion. The capacity of the cell to migrate was examined using CIM plates 24 hours post-transfection with the ASOs and siRNAs targeting the snoRNAs or their host genes’ mature RNA, and the number of migrated cells measured in real time. As indicated in Figure 5A, the knockdown of SNORA19 and its host gene affected cell migration indicating at least in this case that the inhibition of cell migration is caused in part by the expression of EIF3A. In contrast, the knockdown of SNORA81 and not its host gene inhibited cell migration indicating that the effect on migration is specific to SNORA81 (Figure 5B). Similarly, the knockdown of SNORA81 and not SNORA19 inhibited the rate of wound healing in a host gene independent manner (Figure 5C). Indeed, while most cells either completely or partially healed the wound after 48 hours post-scratching, those transfected with ASOs against SNORA81 did not (Figure 5C 81/1 and 81/2 bottom panel). The cell invasion was assessed similarly to the cell migration except a layer of Matrigel coating the upper chamber of the CIM plate permitted to determine the cells invasion rate after the depletion of either the snoRNAs or the host genes. Similarly to the cell migration assay, the knockdown of both SNORA19 and EIF3A inhibited cells’ invasion ability (Figure 5D), while only the knockdown of SNORA81 and not its host gene repressed cells invasion (Figure 5E). We conclude that HGSC-associated snoRNAs may promote cell migration and invasion in a host independent manner at least in the case of SNORA81.

**Figure 5.**
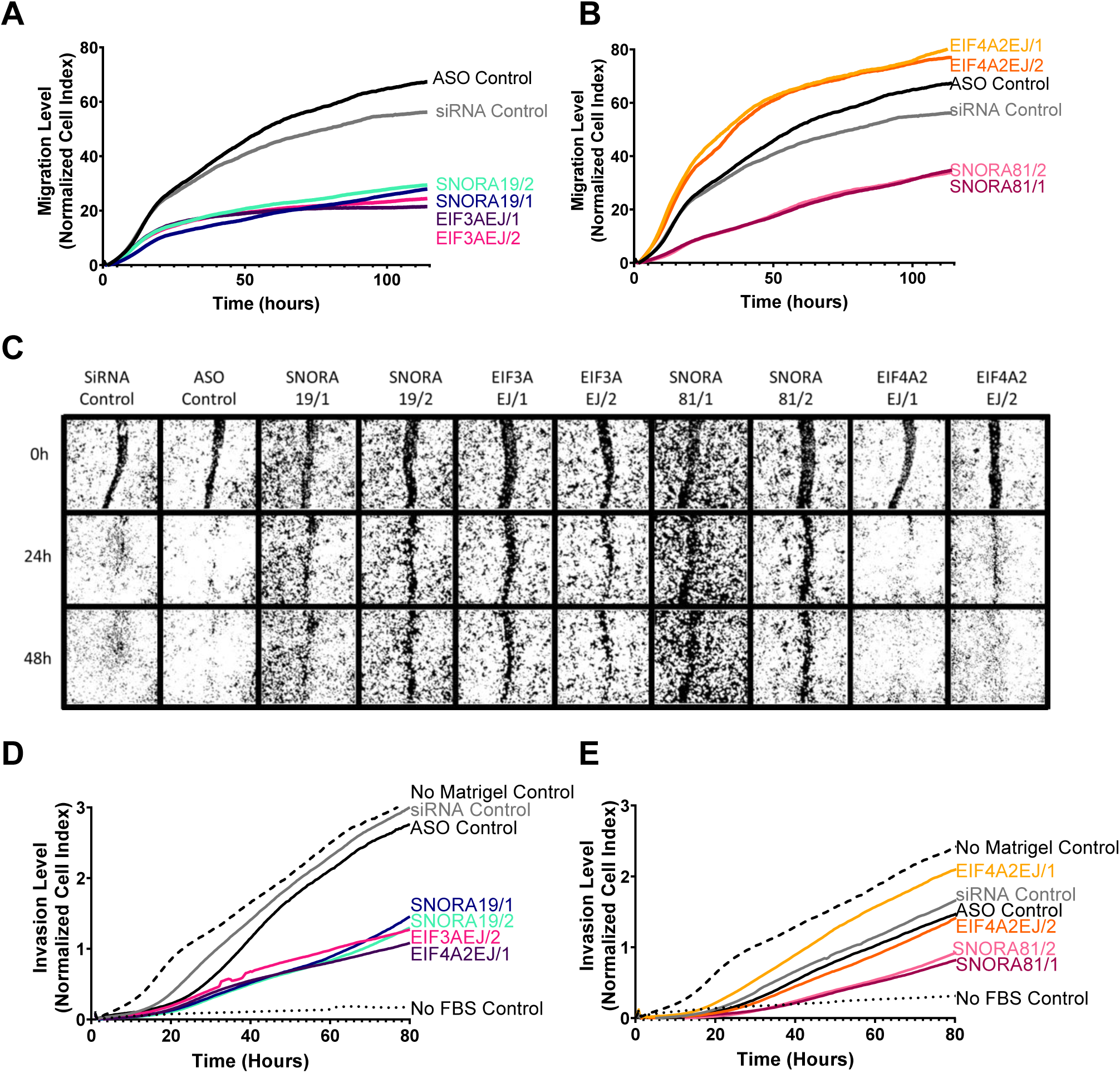
HGSC-associated snoRNAs promote cell migration. **(A & B)** Knockdown of SNORA19 (A) and SNORA81 (B) inhibit cell migration. snoRNA (SNORA19/1, SNORA19/2, SNORA81/1 and SNORA81/2) and their host genes (EIF3AEJ/1, EIF3AEJ/2, EIF4A2EJ/1 and EIF4A2EJ/2) were knocked down in SKOV3ip1 as described in Figure 4B and C and cell migration monitored using Xcelligence RTCA after 24h of transfection. A scrambled ASO (black) and a scrambled siRNA (grey) were used as controls. The y-axis indicate the normalized cell index, which is the average of the three technical replicates. The experiment was carried out in three biological replicates. **(C)** Knockdown of SNORA19 and SNORA81 delays wound healing. Cells were transfected as described in A and wounds introduced 48h after transfection. Healing progress shown after 0, 24, and 48 hours. The experiment was carried out in three biological replicates. The experiment was carried out in three biological replicates. **(D & E)** Knockdown of SNORA19 (D) and SNORA81 (E) inhibit cell invasion. Cells were transfected as described in A and cell invasion was monitored using Xcelligence RTCA CIM plate layered with matrigel 24h after transfection. A scrambled ASO (black) and a scrambled siRNA (grey) were used as controls. No matrigel in upper chamber (dashed line) and no FBS in lower chamber (dotted lines) were used as controls. The y-axis indicate the normalized cell index, which is the average of the three technical replicates. The experiment was carried out in three biological replicates.

## Discussion

In this study, we identified five different snoRNAs that could accurately distinguish between HGSC and SBT with high specificity and accuracy (Figure 3). Interestingly, the knockdown of two examples of these HGSC-associated snoRNAs, SNORA81 and SNORA19, inhibited the proliferation and induced apoptosis of three different model ovarian cancer cell lines consistent with their upregulation in the aggressive HGSC (Figure 4 and Supplementary Figure 5). More importantly, the reduction of these two HGSC-associated snoRNAs inhibited cell migration, wound healing and cell invasion underlining their contribution to the biology of HGSC and tumour aggressiveness. Together the data obtained here suggest a new role for snoRNAs in the promotion of tumour aggressiveness.

Development of cancer, and in particular the increased proliferation rate of cancer cells, is often linked with the dysregulation of translation and ribosome biogenesis (44). Indeed, the increased production of ribosomes is a pre-requisite for increased cell proliferation (45). Accordingly, we hypothesized that the highly invasive and fast growing HGSC cells would also require an overall upregulation of the ribosome production machinery when compared with the much slower growing SBT counterpart. Surprisingly, however, our data indicate that the expression of snoRNAs, key effectors in ribosome biogenesis, is not universally upregulated in HGSC (Figure 2). In fact, the majority of C/D snoRNAs had the tendency to be slightly under expressed in HGSC when compared with SBT. This clearly indicates that the degree of cancer aggressiveness is not strictly linked to the overall induction of snoRNAs. Furthermore, the clear difference in the number of the methylation (C/D) and pseudouridylation (H/ACA) guide snoRNAs that are specifically upregulated in HGSC suggests that these two classes of snoRNA play distinct roles in the development or maintenance of HGSC. Indeed, our work suggests that H/ACA snoRNAs have higher propensity for upregulation in HGSC and are as such more likely to function as positive regulators of cancer cell proliferation and survival. This is consistent with earlier results linking the upregulation of SNORA7B with breast cancer (46) and SNORA42 in lung cancer (47). The link between the upregulation of H/ACA snoRNAs and poor prognosis was also observed in the case of non-small cell lung cancer, where expression of the H/ACA snoRNA binding protein Nop10 was found to drive tumorigenesis (48). However, even within the H/ACA class of snoRNA only 14% were upregulated indicating once more that H/ACA snoRNA expression is not randomly dysregulated in HGSC (Figure 2). Instead, it appears that HGSC requires increased production of a specialized subgroup of H/ACA snoRNAs. The gene specific contribution of H/ACA snoRNAs to tumour aggressiveness is also evident from a previous study showing that the H/ACA SNORA24 may function as a suppressor of hepatocellular carcinoma and its reduced expression is associated with poor prognosis of hepatocellular carcinoma (49). As such, it appears that H/ACA snoRNA may influence carcinogenesis in different ways depending on the identity of the snoRNA and cancer type.

Unlike C/D snoRNAs, which have a high proportion of orphan snoRNAs (not predicted to guide rRNA modifications), most H/ACA snoRNAs guide rRNA modifications and are more tightly linked to changes in translation patterns (16,50,51). Therefore, it is likely that the modulation of the HGSC-associated snoRNAs leads to changes in translation patterns. Indeed, we found that the positions of the predicted modification sites targeted by the HGSC-associated snoRNAs are located predominantly in the 28S rRNA near the peptide exit channel of the ribosome suggesting a role in modulating translation (39,42) and snoRNABase, RRID: SCR_007939) (Supplementary Figure 6A). Notably, except for SNORA19 all four other HGSC-associated snoRNAs have only one predicted modification site, which suggest once more targeted modification of translation. Three of the predicted modification sites to be guided by the H/ACA snoRNAs (positions 3616, 3709 and 4606 of the 28S rRNA) have recently been described as partially modified in HeLa cells suggesting an important regulatory role (52,53) (Supplementary Figure 6B). Consistently, we have shown that SNORA81, which targets the position with the lowest fractional modification in HeLa cells, is enriched in ovarian tissues suggesting that at least in this case, tissue specific variations in snoRNA abundance may contribute to the levels of modification (52–54). The potential of pseudouridylation dependent modification was also shown in different model systems and in cancer cells (55). For example, knockdown of SNORA24 was shown to alter ribosome dynamics leading to increased miscoding and stop codon read through. Furthermore, knockdowns of several H/ACA snoRNAs were shown to alter translation accuracy in yeast (50,56). The exact mechanism by which H/ACA snoRNAs may modify translation will become clearer as more example of cancer associated snoRNA are identified. Meanwhile, the data presented here identify a new subclass of H/ACA snoRNAs that are associated with tumour aggressiveness and demonstrate the requirement for cell proliferation and survival.

## DATA AVAILABILITY

RNA-seq data used in this study are available through the NCBI Gene Expression Omnibus (GEO; https://www.ncbi.nlm.nih.gov/geo) under the accession number GSE181496.

## ACKNOWLEDGEMENTS

We are grateful to Compute Canada for providing state of the art computing facilities to all researchers in Canada.

---

**Supplementary Figure 1 (related to Figures 1-3).**
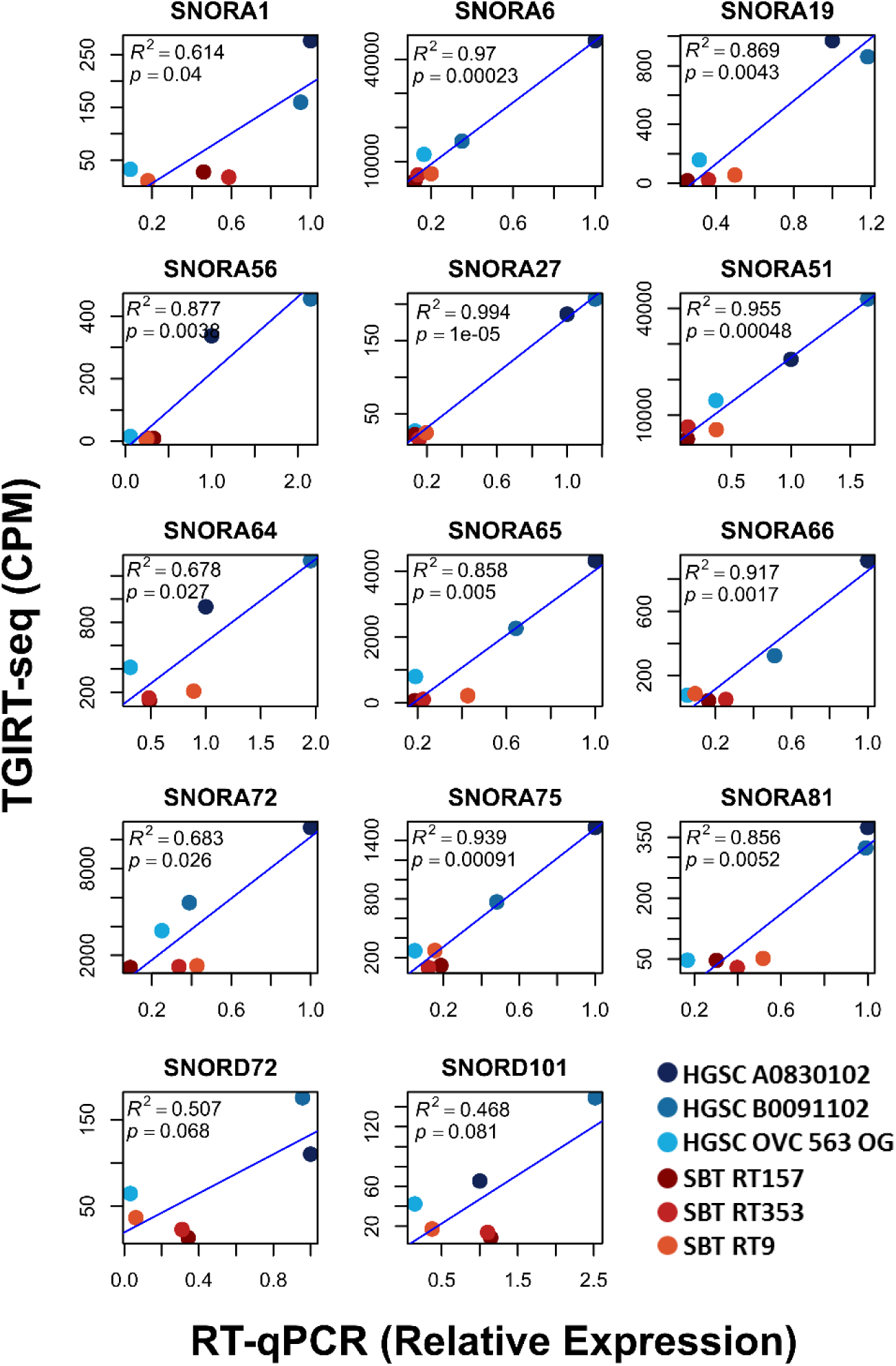
Validation of the TGIRT-seq identified HGSC-associated snoRNA. The abundance of the 14 snoRNAs that were identified by sequencing was examined using RT-qPCR and the results compared to those obtained by sequencing. Each point represents one tissue and those obtained from HGSC and SBT tissues are shown in blue and red shades, respectively. The R squared and its p-value are indicated in the top left corner of each graph.

**Supplementary Figure 2 (related to Figure 3).**
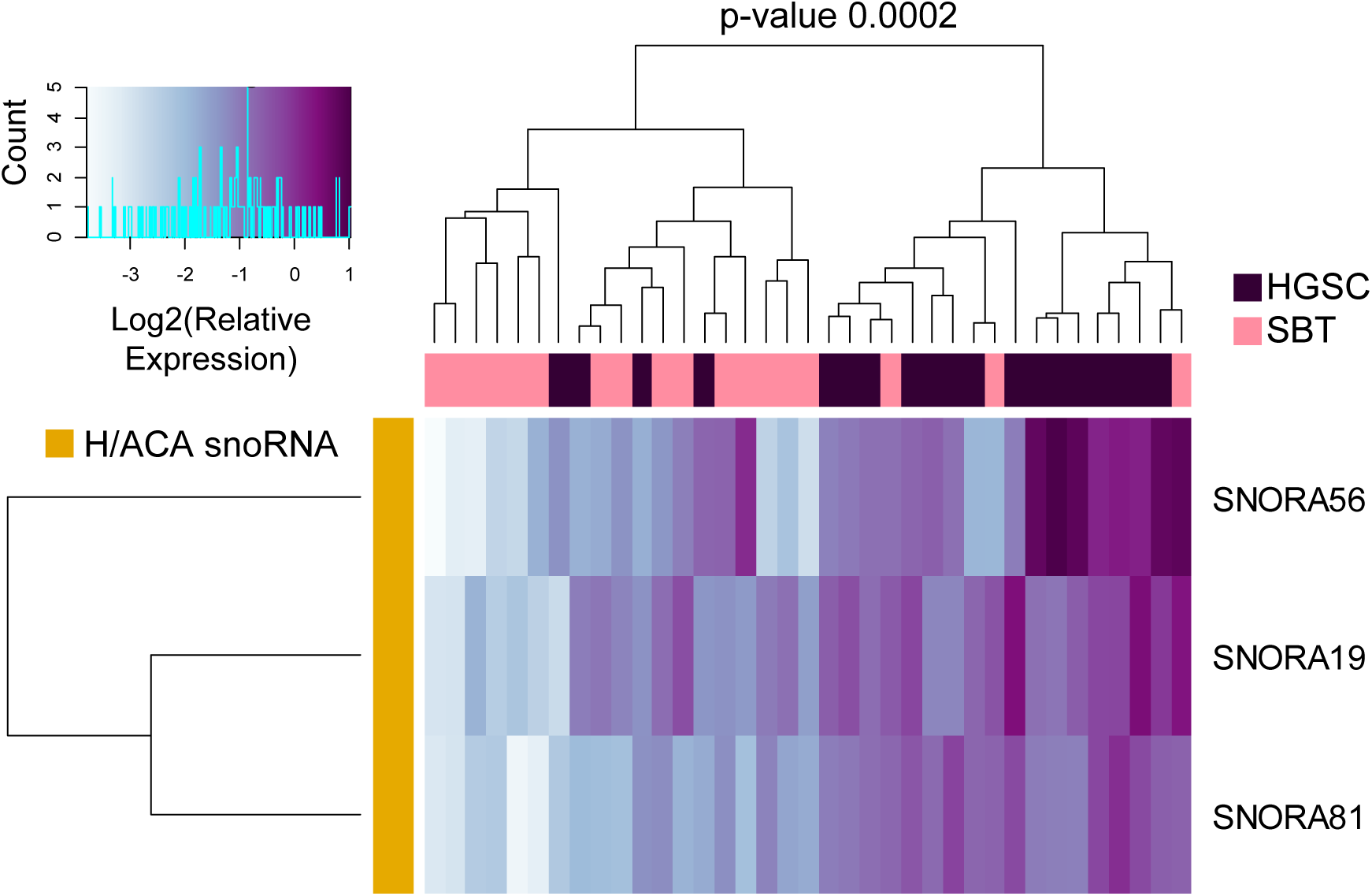
Distribution of the HGSC snoRNA signature in different HGSC and SBT tissues. The relative abundances of the three snoRNAs forming the best HGSC signature identified in Figure 3 were examined in a blind mix of 37 tissues using RT-qPCR. Data were normalized relative to the mean amount of MRPL19 mRNA and a spike-in of E. coli 16S and 23S rRNA and the log2 of the relative expression is presented as a clustered heatmap. The snoRNA names are indicated on the right and the identity of the tissue of origin are shown above. The color key histogram is shown on the top left, where the number of tissue (count) is shown for the color gradient. The Fisher exact test was calculated for the capacity of the snoRNAs to discriminate between HGSC and SBT and shown on top.

**Supplementary Figure 3 (related to Figures 4 and 5).**
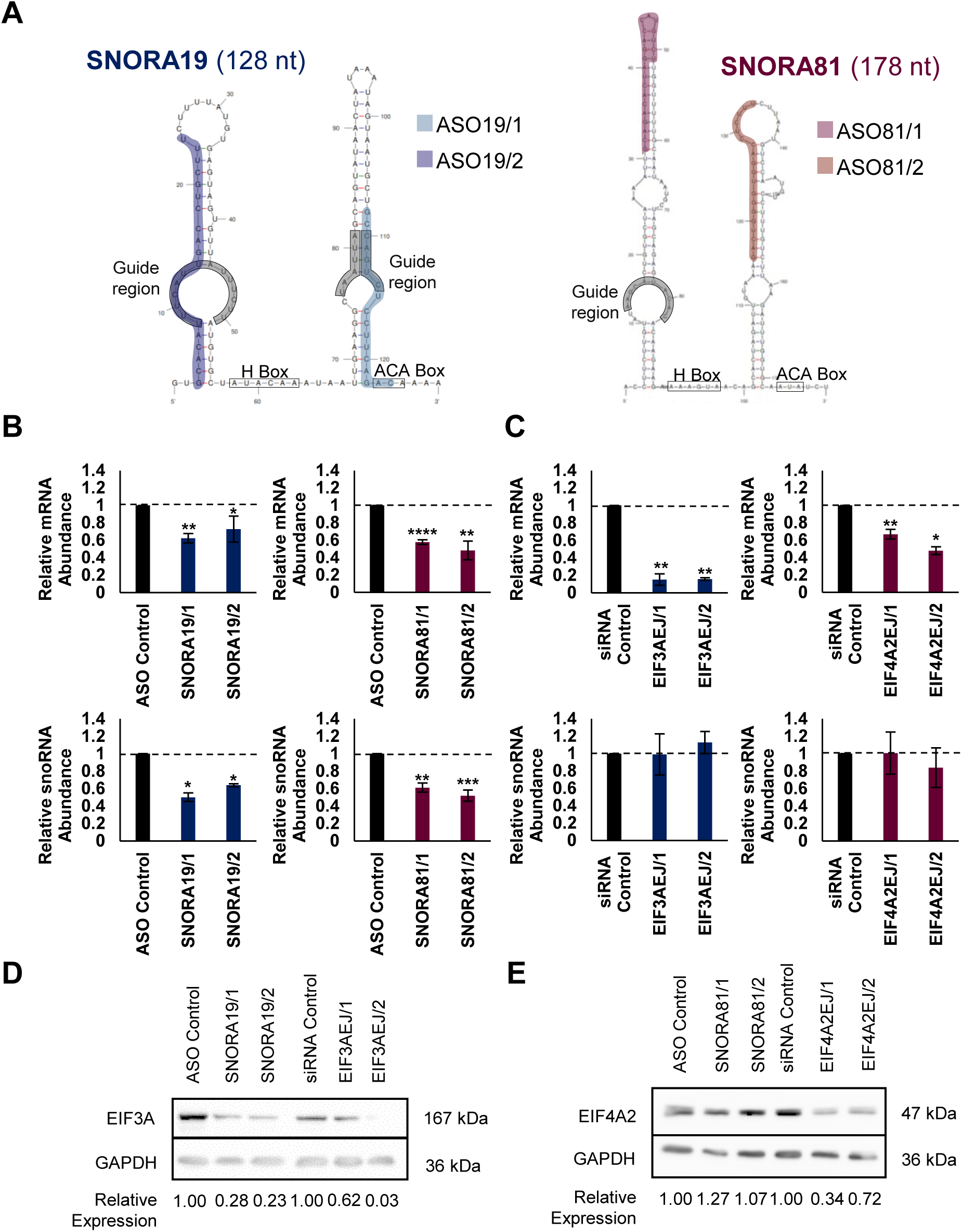
Design and validation of SNORA19 and SNORA81 Knockdowns. **(A)** The structures of SNORA19 and SNORA81 and the positions of the H and ACA boxes, the guide regions and ASOs are highlighted by the grey and blue/purple boxes, respectively. **(B and C)** The relative abundance of EIF3A and EIF4A42 mRNA (upper panels) and the SNORA19 and SNORA81 (lower panels) were determined using RT-qPCR after knockdown using two independent ASOs against each snoRNA (B) and two independent siRNAs against the mature splice junction sequence of each host mRNA (C). The standard deviation obtained from three technical replicates is shown in the form of error bars. *, **, *** and **** indicate p-value (T-test) of 0.04, 0.002, 0.0008, 0.0001, respectively. **(D-E)** Effect of snoRNA and host gene knockdown on the abundance of host protein. The level of EIF3A and EIF4A2 was determined using antibodies specific to each protein before and after knockdown of the snoRNAs and host genes. GAPDH was used as loading control and relative protein levels were quantified and indicated at the bottom. The siRNA and ASO used are indicated on the top and the position of each protein and estimated molecular weight are indicated on each side of the gel.

**Supplementary Figure 4 (related to Figure 4).**
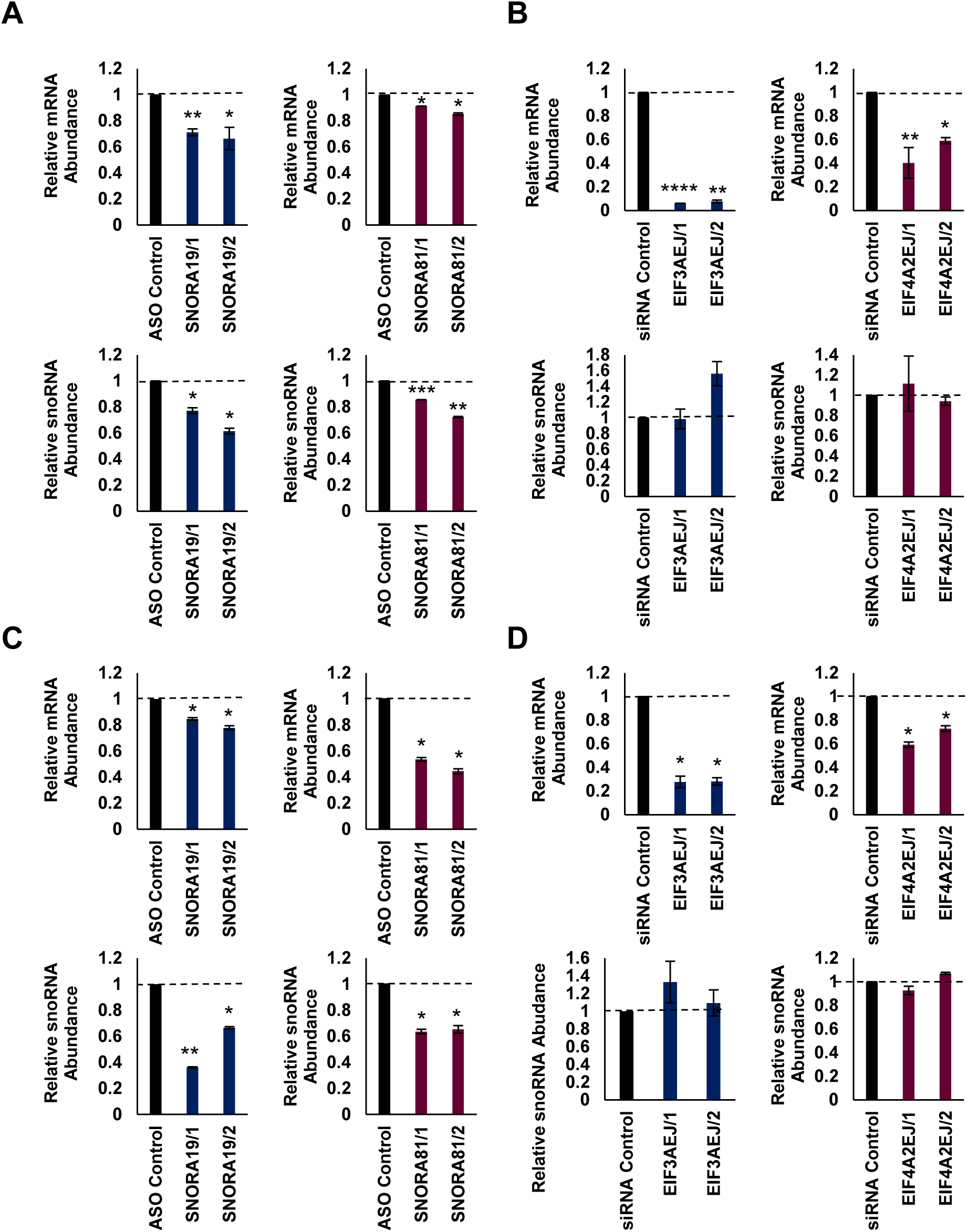
Validation of SNORA19 and SNORA81 knockdowns in different ovarian model cell lines. **(A and B)** Effect of the snoRNAs (A) and their host gene knockdowns (B) on RNA abundance in TOV112D cell line. The abundance of the host mRNA (upper panels) and snoRNA (lower panels) was determined using RT-qPCR before and after knockdown of the snoRNAs (left panels) or their host genes (right panels) as described in Figure S3. *, ** and *** indicate p-value of 0.02, 0.006 and 0.0001 as determined using T-test. **(C and D)** Effect of the snoRNAs (C) and host gene knockdowns (D) on RNA abundance in OVCAR-3 cell line. The effect of knockdown on the snoRNA and host mRNA abundance in the ovarian model cell line OVCAR-3 was determined and illustrated as described in A.

**Supplementary Figure 5 (related to Figure 4).**
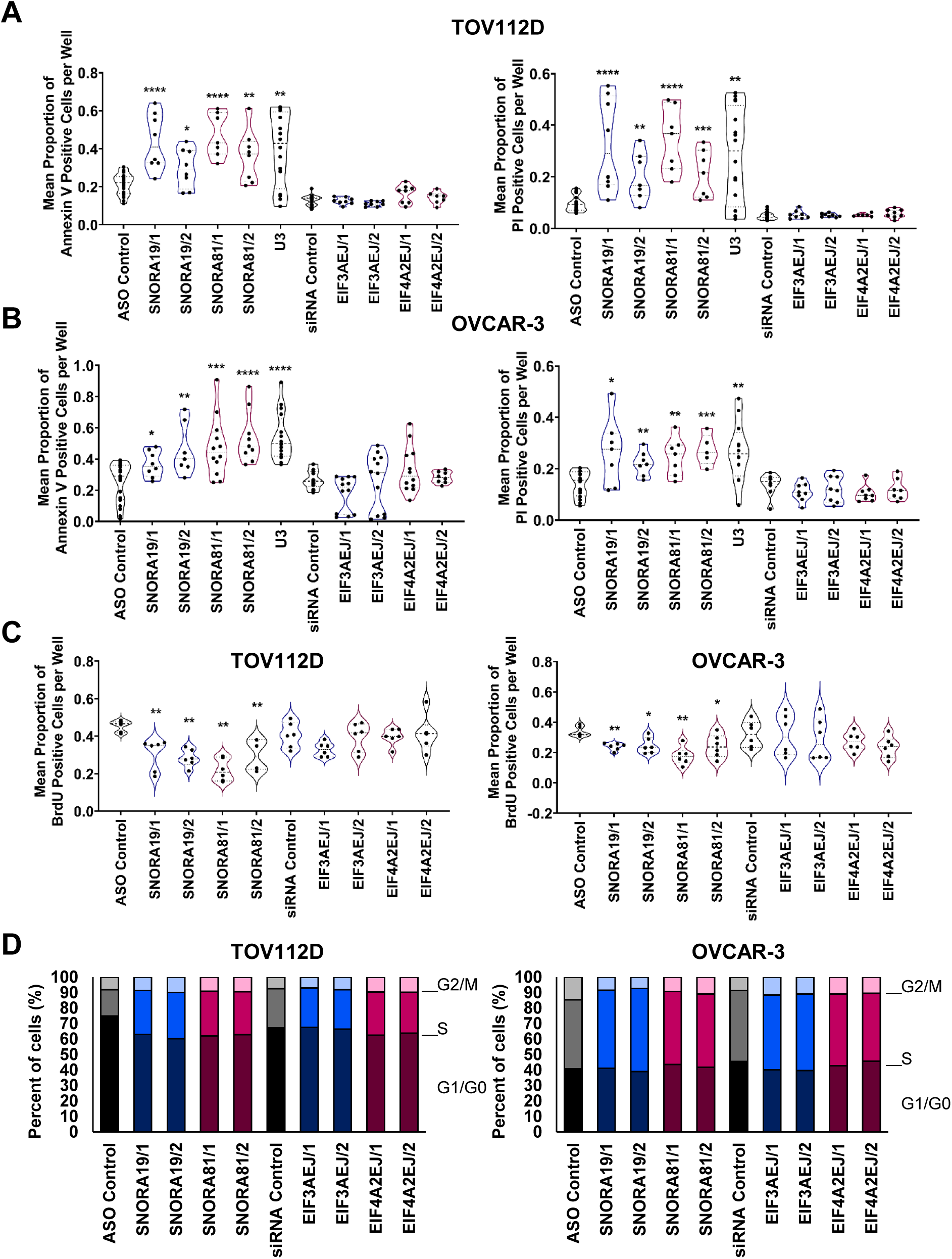
HGSC-associated snoRNAs inhibit apoptosis and necrosis in different ovarian cancer cell lines. **(A)** Knockdowns of SNORA19 and SNORA81 induce apoptosis (left panel) and necrosis (right panel) in TOV112D cell line. The snoRNA and their host genes were knocked down and their effect on apoptosis (annexin V level) and necrosis (propidium iodide level) determined as described in Figure 4. In supplementary Figure 5, stars indicate p-values determined using Mann Whitney test where * are p-value <0.05, ** are p-value >0.01, *** are p-value >0.001 and **** are p-value > 0.0001. The experiment was carried out in three biological replicates with at least four technical replicates. **(B)** Knockdowns of SNORA19 and SNORA81 induce apoptosis (left panel) and necrosis (right panel) in OVCAR-3 cell line. The effect of snoRNA and host gene knockdowns on apoptosis and necrosis of OVCAR-3 was determined and illustrated as described in A. The experiment was carried out in three biological replicates with at least four technical replicates. **(C)** Knockdowns of SNORA19 and SNORA81 inhibit cell proliferation in the TOV112D (right panel) and in the OVCAR-3 (left panel) cell lines. The snoRNA and host gene knockdowns and proliferation assay was performed as described in Figure 4F. The experiment was carried out in three biological replicates with at least four technical replicates. **(D)** Knockdowns of SNORA19 and SNORA81 cause arrest in S phase of the cell cycle in the TOV112D (right panel) and in the OVCAR-3 (left panel) cell lines. The knockdown and cell cycle assay were performed as described in Figure 4G. The experiment was carried out in three biological replicates.

**Supplementary Figure 6.**
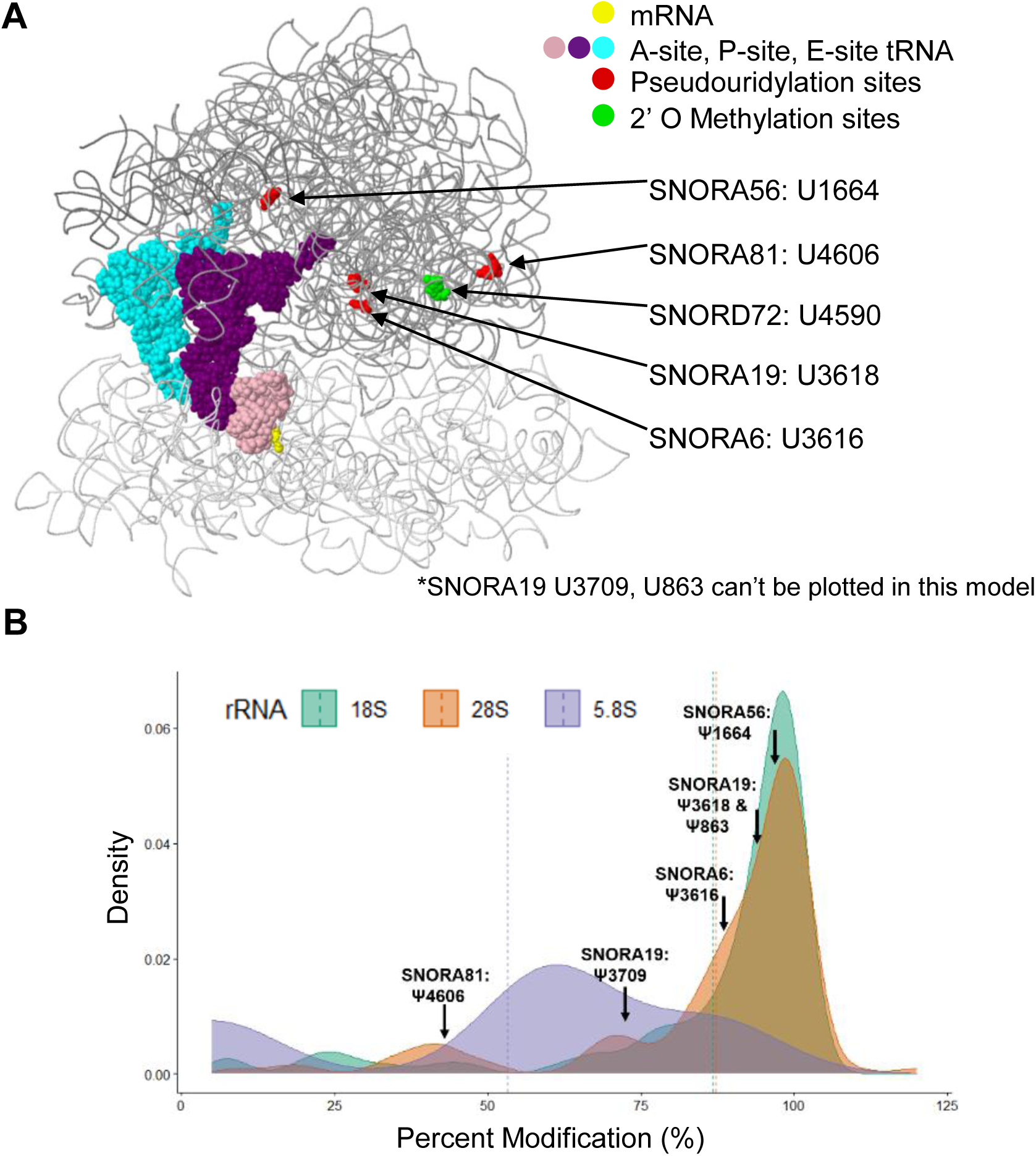
HGSC associated snoRNAs are predicted to guide rRNA modification near the peptide exit channel. **(A)** The 3D structure of the ribosome showing the different predicted rRNA modification of the HGSC associated snoRNAs. Most modifications are located close to the peptide exit channel. Structure was generated using the online database described in Piekna-Przybylska et al (2008) NAR. doi: 10.1093/nar/gkm855. Epub 2007 Oct 18. **(B)** The percent modification of each modification in the respective rRNA. Shown here as a density plot are the specific rRNA modifications predicted to be guided by HGSC associated H/ACA snoRNAs which are all located within the 28S (orange distribution). Data was obtained from Taoka et al (2018) NAR. doi: 10.1093/nar/gky811.

